# Development of KASP markers for the potato virus Y resistance gene *Rychc* using whole-genome resequencing data

**DOI:** 10.1101/2023.12.20.572658

**Authors:** Kenji Asano, Jeffrey B. Endelman

## Abstract

Potato virus Y is the most important potato virus worldwide, affecting tuber yield and quality. The resistance gene *Rychc*, derived from the potato wild relative *Solanum chacoense*, provides broad spectrum and durable resistance to the virus and has been used to develop resistant cultivars. Several DNA markers have been developed and have contributed to the efficient selection of resistant individuals. In this study, we developed Kompetitive Allele Specific PCR markers for *Rychc* using whole-genome resequencing data for a diverse set of 25 PVY susceptible cultivars and a *Rychc*-positive clone. Marker Ry_4099 targets two variants in the 3’-UTR and was able to discriminate all five allele dosages in a tetraploid test population. Marker Ry_3331 targets two variants in Exon 4 and, although it only provides presence/absence information, it discriminates between the two known resistant alleles of *Rychc*. These markers will greatly contribute to efficient development of resistant cultivars.

## Introduction

Potato virus Y (PVY), a species of genus *Potyvirus* in the family Potyviridae, is the most economically important virus of potato (*Solanum tuberosum* L.) worldwide, affecting tuber yield and quality (Adams et al. 2005; Brunt 2001; Karasev and Gray 2013). PVY is mechanically transmitted by more than 40 aphid species in a non-persistent manner (Brunt 2001). Complete control of PVY is difficult even with the application of several management practices, including the use of insecticides and mineral oil, planting certified seed potatoes, sanitizing planting equipment, destroying overwintering sources of PVY inoculum, using crop borders, and timely rogueing of volunteer potato plants and weed reservoirs of aphids and virus (MacKenzie et al. 2014). Resistant cultivars provide the most effective and durable control of PVY.

Two types of resistance to PVY are known in potato: hypersensitive resistance (HR) and extreme resistance (ER). HR is conferred by *Ny* genes and results in necrotic lesions inhibiting viral spread from cell-to-cell and through the vascular system (Valkonen 2015; Valkonen et al. 1996). HR genes often confer strain specific resistance and are not effective under some conditions such as high temperature (Jones 1990; Valkonen 1997). By contrast, ER is conferred by *Ry* genes and show broad spectrum and durable resistance. With ER, the virus is either undetectable or detectable only with highly sensitive techniques (Ohki et al. 2018; Valkonen et al. 1996). To date, three genes that confer ER to PVY have been known from three different sources: (1) *Rysto* from *S. stoloniferum* Schltdl. et Bouché (Cockerham 1943), (2) *Ryadg* from *S. tuberosum* L. subsp. *andigena* Hawkes (Munoz et al. 1975), and (3) *Rychc* from *S. chacoense* Bitt. (Asama et al. 1982). Although weakening of resistance by the *Rychc* gene has also been reported under high temperature (Ohki et al. 2018), there have been no reports of PVY strains that have overcome ER to date, so the introduction of *Ry* genes is considered an effective strategy for durable resistance against PVY.

Recently, *Rychc* was cloned from different sources by independent studies (Akai et al. 2023; Li et al. 2022). PVY resistance derived from *S. chacoense* was first introduced into a Japanese commercial cultivar Konafubuki by Asama et al. (1982). Subsequently the gene was genetically identified as a single dominant gene and named *Rychc* (Hosaka et al. 2001). The origin of the *Rychc* gene in Konafubuki can be traced back to *S. chacoense* W84. On the other hand, Li et al. (2022) cloned the resistance gene from another source, *S. chacoense* 40-3. Sequence comparison revealed that there are several differences, leading to several amino acid changes between these alleles (Akai et al. 2023). Therefore, we renamed *Rychc* derived from Konafubuki and *S. chacoense* 40-3 as *Rychc-1* and *Rychc-2*, respectively.

Kompetitive Allele Specific PCR (KASP) is a simplified fluorescence-based methodology for single nucleotide polymorphism (SNP) or insertion/deletion genotyping assays in which the DNA sample is amplified using a thermal cycler and allele-specific primers. KASP allows breeders to screen genotypes in a high-throughput and cost-effective manner while avoiding the limitations of post-PCR handling (Semagn et al. 2014). Based on the fluorescence ratio for the two alleles, all three genotypes in a diploid are readily distinguished, and the different heterozygote dosages in a polyploid can sometimes be inferred (Kante et al. 2021; Sorensen et al. 2023; Uitdewilligen et al. 2013). To date, KASP markers for *Rysto* and *Ryadg* have been developed (Kante et al. 2021), but not for *Rychc*. Our objective was to develop KASP markers for *Rychc*.

## Materials and methods

### Identification of variants for KASP marker design

Sequencing data for 23 North American (Hardigan et al. 2017) and two Japanese cultivars (Yamakawa et al. 2021) were downloaded from the European Nucleotide Archive (accession PRJNA378971 for North American and PRJDB10795 for Japanese cultivars, Supplemental Table 1) to use as *Rychc*-negative samples. In addition, Illumina HiSeq 2x100 bp data of 98H20-5, a diploid clone heterozygous for *Rychc-1* (Sato et al. 2006), was generated by Eurofins Genomics (Chiyoda, Tokyo, Japan). Reads were aligned to the *Rychc-1* genomic sequence (GeneBank ID: LC726345) using BWA-MEM version 0.7.17 (Li 2013). SAMtools version 1.13 (Danecek et al. 2021) was used to remove unmapped reads and PCR duplicates. The software IGV (Robinson et al. 2011) was used to visualize the alignments. A VCF file was generated using FreeBayes v1.3.5 (Garrison and Marth 2012) and processed using the vcfR package (Knaus and Grunwald 2017).

Homologous gene fragments from *S. chacoense* accessions identified by Li et al. (2022) were downloaded from the NCBI database and aligned using MAFFT with default parameters in UGENE (Okonechnikov et al. 2012).

### Plant materials

KASP makers were tested on diploid and tetraploid populations for which *Rychc* segregation was expected. The diploid population consisted of 48 F_2_ progeny from the cross DM1-3 x M6 (Endelman and Jansky 2016), chosen because M6 contains *Rychc-2* (Li et al. 2022) . The population was assayed using 2K targeted genotyping-by-sequencing (DArTag by Diversity Arrays Technology, Bruce, ACT Australia). PolyOrigin (Zheng et al. 2021) was used for linkage analysis of the F_2_ (S_1_) population to reconstruct progeny in terms of parental haplotypes and predict the dosage of the *Rychc*-bearing M6 haplotype, based on reference genome DM1-3 v6.1 coordinates (Pham et al. 2020). The tetraploid population consisted of 88 clones derived from 37 crosses of the UW-Madison potato breeding program, of which 51 were candidates for *Rychc* because they were grandchildren of AW07791-2rus. AW07791-2rus is a male fertile breeding line with maternal parent PALB0303-1, a known carrier of *Rychc* (Elison et al. 2021). AW07791-2rus tested positive for *Rychc* based on SCAR markers RY1648 and MG64-17, which were designed from the gene sequence and show no discrepancy with the resistance phenotype (Akai et al. 2023; Li et al. 2022). A breeding line Saikai 35 contains *Rychc-1* as a descendant of Konafubuki (Mori et al. 2012) and was used to develop a KASP marker that discriminates between the alleles *Rychc-1* and *Rychc-2*.

### Experimental Protocols

Primers for the SCAR markers were obtained from Integrated DNA Technologies (Coralville, Iowa, USA). DNA was extracted from lyophilized leaf or tuber tissue with KingFisher Duo (Thermo Fisher Scientific, Waltham, Massachusetts, USA) using the NucleoMag^®^ Plant kit (MACHEREY-NAGEL, Düren, German), and concentration was adjusted to 10 ng/μl before the PCR and KASP assays. PCR reactions were conducted in 10 μL volume comprising 5 μL Promega 2x PCR Master Mix (Promega, Madison, Wisconsin, USA), 10 pmol of forward and reverse primers, and 2 μL of template DNA. Thermal cycling was run on a Bio-Rad C1000 Touch 96 well thermal cycler (Bio Rad, Hercules, California, USA). The PCR conditions for RY1648 consisted of one cycle of 5 min at 94 □; followed by 35 cycles of 94 □for 30 s, 55□for 30 s, and 72 □for 1 min; and one final cycle of 5 min at 72□. The PCR conditions for MG64-17 consisted of one cycle of 3 min at 94 □; followed by 35 cycles of 94 □for 30 s, 55□for 30 s, and 72 □for 30 s; and one final cycle of 10 min at 72□. 2.0% agarose gel was made with 0.5x TBE and 0.01% of SYBR^®^ Safe DNA gel stain (Thermo Fisher Scientific). 2 μL of the 6x loading dye was added to 10 μL of the PCR product and loaded into the gel. The gel was electrophoresed with the VWR^®^ Real Time Electrophoresis System (Avantor, Radnor, Pennsylvania, USA) in a 0.5x TBE bath for 40 min at 100 V. The banding results were visualized with a blue LED transilluminator. Sequences of SCAR markers used in this study are listed Supplemental Table 2.

The KASP markers were designed by LGC Biosearch Technologies (Teddington, UK) based on the target SNPs and flanking sequences. For ordering, use Project ID 1657.010 for marker Ry_3331, and Project ID 1657.011 for Ry_4099. The assay was conducted in a 10 μL reaction volume, comprising 5 μL of 2x KASP master mix, 0.14 μL of assay mix, and 5 μL of template DNA. Thermal cycling was run on a Bio-Rad C1000 Touch 96 well thermal cycler, using the standard 61-55 °C touchdown protocol for Ry_4099 and the 2-step 57□protocol for Ry_3331. The standard 61-55 °C touchdown protocol is 94 ºC for 15 min; 10 cycles of 94 ºC for 20 s and 61 ºC for 1 min with a decrease of 0.6 ºC every cycle; 26 cycles of 94 ºC for 20 s and 55 ºC for 1 min; a final plate reading step at 30 ºC for 1 min. The 2-step 57□protocol is 94 ºC for 15 min; 36 cycles of 94 □for 20 s and 57 □for 1 min; a final plate reading step at 30 □for 1 min. Clusters were defined based on the slope of their regression line according to the ratio of HEX: FAM fluorescence.

### Results

To identify variants specific to *Rychc*, Illumina short reads for 25 cultivars without extreme resistance to PVY were aligned to the *Rychc-1* genomic sequence. As an additional positive control, approximately 50X whole-genome sequence data for 98H20-5, a diploid clone heterozygous for *Rychc-1*, was included. As seen in other studies (Sorensen et al. 2023), the alignment depth in exonic regions was much higher than the expected coverage, indicating the presence of reads from homologous genes (Figure 1, Supplemental Table 3). Thus, only variants in the intron and UTR regions were considered further. Five variants met our three criteria: (1) identical REF alleles in *Rychc-1* and *Rychc-2*; (2) homozygous ALT for all 25 susceptible cultivars; and (3) 98H20-5 was HET. Two variants were discarded during the design phase, and one was tested but failed (Table 1).

**Figure 1.**
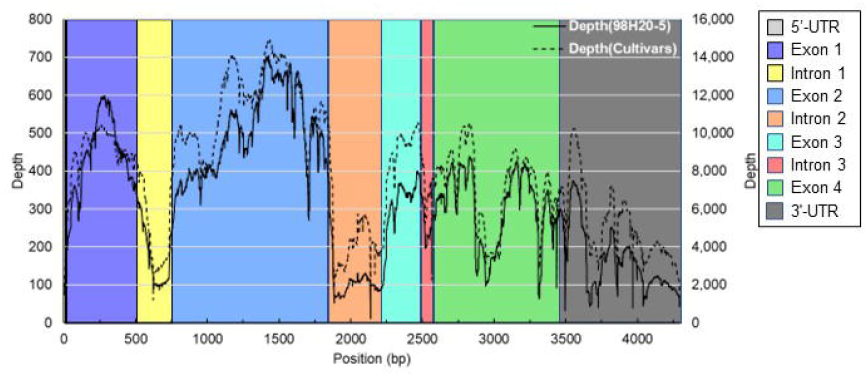
Read depth from the whole genome sequencing of 98H20-5 and 25 cultivars. Solid and dotted lines represent the read depth of 98H20-5 (left Y axis) and total read depth from 25 cultivars (right Y axis), respectively. 5’- and 3’-UTRs, exon, and intron regions are indicated by different colors.

**Table 1.**
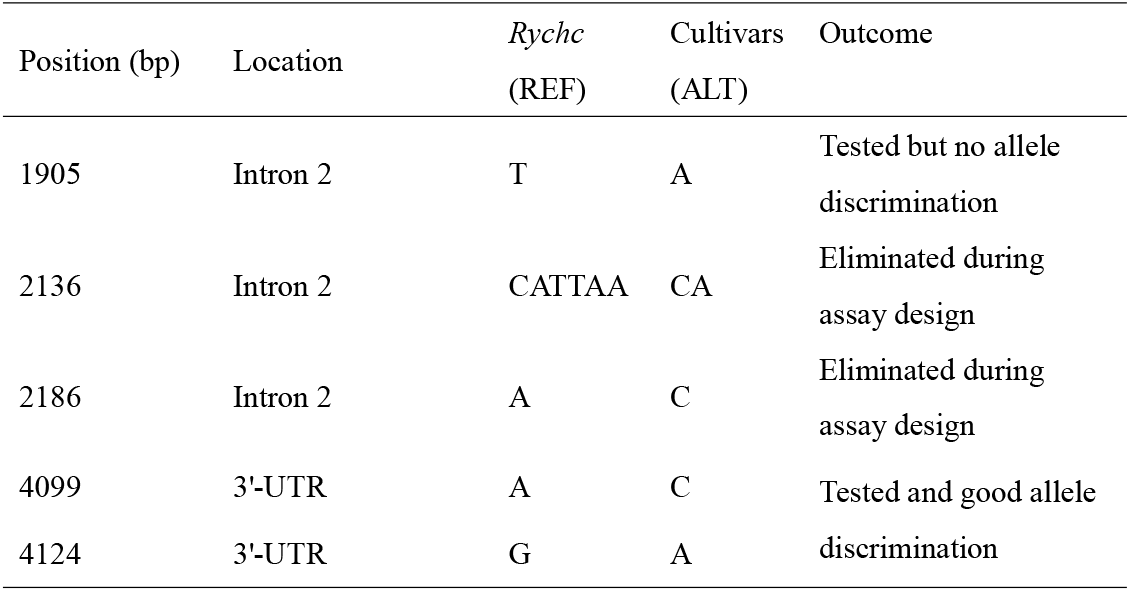
Candidate variants for the KASP markers. Position (bp) is from the transcription start site of *Rychc-1*.

The last two candidate variants, located in the 3’-UTR at 4099 bp [A/C] and 4124 bp [G/A], were combined to design marker Ry_4099. In a test set of 88 tetraploid samples, Ry_4099 generated two clusters in perfect agreement with the previously validated SCAR markers for *Rychc* (Figure 2a). In a F_2_ diploid population, three distinct clusters were observed (Figure 2b). Once again, the predicted presence/absence of *Rychc* was in perfect agreement with the SCAR marker, and parental haplotype analysis using genome-wide markers validated the discrimination between *RR* and *Rr* genotypes, i.e., the codominant behavior of Ry_4099. To assess the potential for allele dosage discrimination in a tetraploid population, a triplex genotype (*RRRr*) was simulated by 1:1 mixture of *Rr* and *RR* DNA samples. In a test population with all five possible *R*:*r* ratios (1:0, 3:1, 1:1, 1:3, 0:1), five distinct clusters were observed (Figure 3).

**Figure 2.**
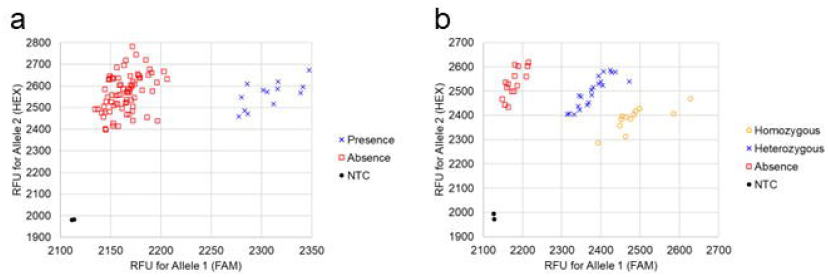
Genotyping result with KASP marker Ry_4409. a and b: Results for tetraploid (a) and diploid (b) population. X- and Y-axes show relative fluorescence units, RFU, for FAM (Allele1) and HEX (Allele2). NTC = no template control.

**Figure 3.**
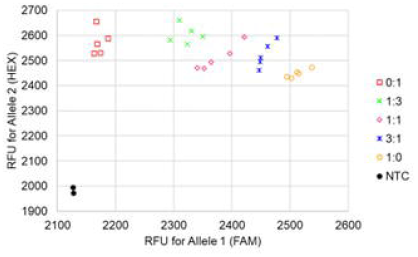
Result of allele dosage discrimination with Ry_4099. X- and Y-axes show relative fluorescence units, RFU, for FAM (Allele1) and HEX (Allele2). 1:0=absence, 3:1=simplex in tetraploid, 1:1=heterozygous in diploid, 1:3=1:1 mixture of *Rr* and *RR*, 0:1= homozygous for *Rychc* in diploid, NTC = no template control.

Although no functional differences between *Rychc-1* and *Rychc-2* have been established yet, there may be a future need to discriminate between these alleles. By comparing their genomic sequences, candidate variants were evaluated for KASP marker design, with the additional constraint that *Rychc*-negative samples behave as null alleles. KASP marker Ry_3331 exploits two variants in Exon4: an *Rychc*-specific variant at 3308 bp [AG/TC] and a SNP at 3331 bp [C/T] that differentiates the two *Rychc* alleles (Figure 4). Applying Ry_3331 to our diploid and tetraploid test populations, separate *Rychc*-positive clusters were observed for the two ploidies, along with a *Rychc*-negative cluster at the origin of the fluorescence scatter plot (Figure 5). The tetraploid test samples grouped with the *Rychc-1* control sample, Saikai 35. There were no discrepancies between the predictions of Ry_3331 and the SCAR markers (Figure 5).

**Figure 4.**
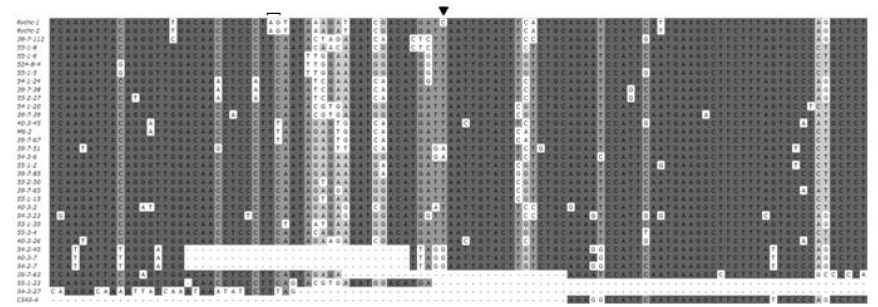
Comparison of sequences used for KASP design. *Rychc-1, Rychc-2* and homologous genes were aligned. Sequence region submitted to LGC and corresponding regions of the homologous genes are shown. Closing bracket and inverted triangle indicate variants used for KASP development.

**Figure 5.**
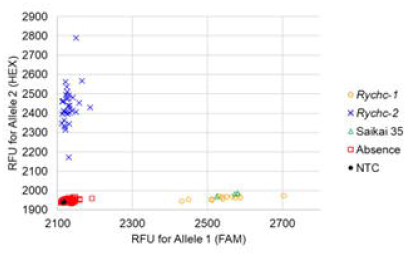
Genotyping result with KASP marker Ry_3331 discriminating *Rychc-1* and *Rychc-2*. Saikai 35 was included with three replicates. X- and Y-axes show relative fluorescence units, RFU, for FAM (Allele1) and HEX (Allele2). NTC = no template control.

## Discussion

With the recent advance in sequencing technologies, several methods have been proposed to efficiently develop DNA markers using whole genome sequence of bulked DNA (Chen et al. 2019; Chen et al. 2018; Clot et al. 2020; Meade et al. 2020; Strachan et al. 2019; Yamakawa et al. 2021). When only a few biparental populations are used to identify SNPs linked to the target genes, these SNPs may not be effective in other genetic backgrounds. Combining these methods with confirmation of the variants using whole-genome resequencing data of diverse cultivars can identify more robust markers.

Ry_4099 is a codominant marker discriminating all possible *Rychc* genotypes. In recent years, there has been a great deal of interest in diploid potato breeding and the production of diploid hybrid potato cultivars (Bethke et al. 2022; Jansky et al. 2016; Lindhout et al. 2011; Zhang et al. 2021). Once inbred lines are developed, useful genes can be rapidly incorporated through marker-assisted backcrossing (Bradshaw 2022; Su et al. 2020). In tetraploid populations, breeding materials with multiple copies of useful genes are valuable for improving breeding efficiency (Andrade et al. 2009; Kaushik et al. 2013; Mori et al. 2015). Ry_4099 will greatly contribute to the rapid selection of *Rychc* homozygous individuals in diploid breeding programs and multiplex genotypes for *Rychc* in tetraploid breeding.

To date, no differences in strain specificity or resistance responses have been reported between *Rychc-1* and *Rychc-2*. However, there are several sequence differences leading to amino acid exchanges, including in the TIR, NBS, and LRR domains (Akai et al. 2023). These domains are thought to contribute to pathogen recognition specificity and control of the defense response (DeYoung and Innes 2006; Duxbury et al. 2021; Jones et al. 2016), so there is the possibility of functional differences now or in the future, particularly if *Rychc* becomes widely deployed in commercial production. At that time, it could be important to discriminate between *Rychc-1* and *Rychc-2*.

## Acknowledgements

We thank Dr. Kazuyoshi Hosaka, Obihiro University of Agriculture and Veterinary Medicine, and Dr. Xingkui Cai, Huazhong Agricultural University, for providing 98H20-5 and sequence information of *Rychc-2*, respectively. We also thank Dr. Kotaro Akai, Hokkaido Agricultural Research Center, National Agricultural Research Organization, and Grace Christensen, University of Wisconsin-Madison, for helpful suggestions on the research. This study was supported by grants from the Ministry of Agriculture, Forestry, and Fisheries of Japan (Genomics-Based Technology for Agricultural Improvement [SFC3002]), and USDA NIFA Award 2021-34141-35447.

## Author Contribution

KA designed the study, conducted the experiments, wrote the original draft, and edited the manuscript. JBE contributed methods, data, and germplasm, supervised the project, reviewed, and edited the manuscript.

## Data Availability

Whole-genome sequence data for 98H20-5 is available from the DDBJ Sequence Read Archive (DRA) under BioProject ID PRJDB16925.

