## Supplemental Tables for "Development of KASP markers for the potato virus Y resistance gene *Rychc* using whole-genome resequencing data"

Supplement materials

Supplemental Table 1 Sequence data used for this study.

| Sample Accession | Experiment Accession | Run Accession | Name/Accession | Reference |
| --- | --- | --- | --- | --- |
| SAMD00655053 | DRX495505 | DRR511619 | 98H20-5 | this study |
| SAMN06564743 | SRX2645993 | SRR5349577 | Snowden | Hardigan *et al.* (2017) |
| SAMN06564623 | SRX2645994 | SRR5349578 | Atlantic | Hardigan *et al.* (2017) |
| SAMN06564742 | SRX2645997 | SRR5349579 | Kalkaska | Hardigan *et al.* (2017) |
| SAMN06564741 | SRX2645998 | SRR5349580 | Missaukee | Hardigan *et al.* (2017) |
| SAMN06564740 | SRX2645999 | SRR5349581 | Russet Norkotah | Hardigan *et al.* (2017) |
| SAMN06564738 | SRX2646001 | SRR5349583 | Yukon Gold | Hardigan *et al.* (2017) |
| SAMN06564737 | SRX2646002 | SRR5349584 | Spunta | Hardigan *et al.* (2017) |
| SAMN06564736 | SRX2646003 | SRR5349585 | Sierra Gold | Hardigan *et al.* (2017) |
| SAMN06564735 | SRX2646004 | SRR5349586 | Shepody | Hardigan *et al.* (2017) |
| SAMN06564734 | SRX2646005 | SRR5349587 | Russet Burbank | Hardigan *et al.* (2017) |
| SAMN06564733 | SRX2646006 | SRR5349588 | Rio Grande Russet | Hardigan *et al.* (2017) |
| SAMN06564732 | SRX2646007 | SRR5349589 | Purple Majesty | Hardigan *et al.* (2017) |
| SAMN06564731 | SRX2646008 | SRR5349590 | Premier Russet | Hardigan *et al.* (2017) |
| SAMN06564730 | SRX2646009 | SRR5349591 | Norland | Hardigan *et al.* (2017) |
| SAMN06564729 | SRX2646010 | SRR5349592 | Mountain Rose | Hardigan *et al.* (2017) |
| SAMN06564728 | SRX2646011 | SRR5349593 | Kennebec | Hardigan *et al.* (2017) |
| SAMN06564727 | SRX2646012 | SRR5349594 | Katahdin | Hardigan *et al.* (2017) |
| SAMN06564726 | SRX2646013 | SRR5349595 | Irish Cobbler | Hardigan *et al.* (2017) |
| SAMN06564725 | SRX2646014 | SRR5349596 | Garnet Chili | Hardigan *et al.* (2017) |
| SAMN06564724 | SRX2646015 | SRR5349597 | Early Rose | Hardigan *et al.* (2017) |
| SAMN06564723 | SRX2646016 | SRR5349598 | Dakota Diamond | Hardigan *et al.* (2017) |
| SAMN06564722 | SRX2646017 | SRR5349599 | Burbank | Hardigan *et al.* (2017) |
| SAMN06564681 | SRX2646060 | SRR5349641 | Superior | Hardigan *et al.* (2017) |
| SAMD00258699 | DRX245172 | DRR255454 | Sayaka | Yamakawa *et al.* (2021) |
| SAMD00258700 | DRX245173 | DRR255455 | Hokkaikogane | Yamakawa *et al.* (2021) |

Supplemental Table 2 Sequence of SCAR markers used for this study.

| DNA marker | Primer name | Primer sequence (5'→3') | Expected size (bp) | Reference |
| --- | --- | --- | --- | --- |
| RY1648 | 1648F24 | ACAACCTCCCTAGTATAAAGATGATCGACATGATC | 594 | Akai *et al* (2023) |
|  | 1648R22 | GTATAACAGATGGATCCCTATCTTCTTTACAAC |  |  |
| MG64-17 | MG64-17-F | TAAGTAAGAAACCTACTTATTCTCCG | 882 | Li *et al* (2022) |
|  | MG64-17-R | GTTTGACAACCTCCCTAGTATAAA |  |  |

Supplemental Table 3 Average read depth in each region from whole genome re-sequence of 98H20-5 and sum of 25 cultivars.

|  | 5'-UTR | Exon1 | Intron1 | Exon2 | Intron2 | Exon3 | Intron3 | Exon4 | 3'-UTR |
| --- | --- | --- | --- | --- | --- | --- | --- | --- | --- |
| 98H20-5 | 77 | 435 | 179 | 496 | 115 | 304 | 261 | 303 | 167 |
| Cultivars | 2,097 | 8,958 | 4,864 | 11,406 | 4,435 | 8,677 | 7,250 | 7,134 | 5,216 |
